# Generation of PRKN and PINK1-KO and double KO cell lines from healthy induced pluripotent stem cells using CRISPR/Cas9 editing

**DOI:** 10.1101/2022.02.25.482014

**Authors:** Carol X.-Q. Chen, Zhipeng You, Narges Abdian, Julien Sirois, Irina Shlaifer, Mahdieh Tabatabaei, Marie-Noëlle Boivin, Lydiane Gaborieau, Jason Karamchandani, Lenore K. Beitel, Edward A. Fon, Thomas M. Durcan

**Affiliations:** The Neuro’s Early Drug Discovery Unit (EDDU), McGill University, 3801 University Street, Montreal, Quebec, Canada H3A 2B4; C-BIG Biorepository (C-BIG), Montreal Neurological Institute-Hospital, McGill University, Montreal, Quebec, Canada H3A 2B4; McGill Parkinson Program and Neurodegenerative Diseases Group, Montreal Neurological Institute, Department of Neurology and Neurosurgery, Montreal, Quebec, Canada H3A 2B4

## Abstract

Autosomal recessive mutations in either *PRKN* or *PINK1* are associated with early-onset Parkinson’s disease. The corresponding proteins, PRKN, an E3 ubiquitin ligase, and the mitochondrial serine/threonine-protein kinase PINK1 play a role in mitochondrial quality control. Using CRISPR/CAS9 technology we generated three human iPSC lines from the well characterized AIW002-02 control line. These isogenic iPSCs contain homozygous knockouts of *PRKN* (PRKN-KO, CBIGi001-A-1), *PINK1* (PINK1-KO, CBIGi001-A-2) or both *PINK1* and *PRKN* (PINK1-KO/PRKN-KO, CBIGi001-A-3). The knockout lines display normal karyotypes, express pluripotency markers and upon differentiation into relevant brain cells or midbrain organoids may be valuable tools to model Parkinson’s disease.

## 1. Resource Table

**1. Resource Table:**
|  |  |
| --- | --- |
| Unique stem cell lines identifier | CBIGi001-A-1<br>CBIGi001-A-2<br>CBIGi001-A-3 |
| Alternative name(s) of stem cell lines | PRKN-KO/AIW002-02 (PRKN-KO)<br>PINK1-KO/AIW002-02 (PINK1-KO)<br>PINK1-KO/PRKN-KO/AIW002-02 (PINK1-KO/PRKN-KO) |
| Institution | The Montreal Neurological Institute (The Neuro) |
| Contact information of the reported cell line distributor | Thomas Durcan, PhD; |
| Type of cell lines | iPSC |
| Origin | Human |
| Additional origin info | Age: 37<br>Sex: Male<br>Ethnicity: Caucasian |
| Cell Source | AIW002-02 (CBIGi001-A) |
| Clonality | Clonal |
| Method of reprogramming | Parental AIW002-02 iPSCs were derived from PBMCs of a healthy male using a four factor (Klf4, Oct3/4, Sox2, cMyc) Sendai virus-based system |
| Multiline rationale | Isogenic genome-edited disease cell lines |
| Evidence of the reprogramming transgene loss (including genomic copy if applicable) | N/A |
| Cell culture system used | mTESR1 medium on Matrigel coated plates |
| Type of Genetic Modification | Homozygous gene knockout |
| Associated disease | Parkinson's disease |
| Gene/locus | *PRKN* (6q25.2-q27) (77 bp deletion in Exon 2)<br>*PINK1* (1p35-p36) (17 bp deletion/18 bp "Stop-tag" insertion)<br>*PINK1/PRKN* (PRKN-KO: 77 bp deletion in exon 2)/*PINK1*-KO: 5 bp deletion in Exon 1) |
| Method of modification/site-specific nuclease used | CRISPR/Cas9-mediated gene knockout |
| Site-specific nuclease (SSN) delivery method | PRKN-KO: Plasmid transfection<br>*PINK1*-KO and *PINK1*-KO/*PRKN*-KO: Nucleofection |
| All genetic material introduced into the cells | PRKN-KO: CRISPR/Cas9 plasmid |
| Analysis of the nuclease-targeted allele status | Sequencing of targeted alleles |
| Method of the off-target nuclease activity surveillance | Targeted PCR/sequencing |
| Name of transgene | N/A |
| Eukaryotic selective agent resistance (including inducible/gene expressing cell-specific) | PRKN-KO: puromycin |
| Inducible/constitutive system details | N/A |
| Date archived/stock date | 2021-01-22 |
| Cell line repository/bank | https://hpscreg.eu/cell-line/CBIGi001-A-1<br>https://hpscreg.eu/cell-line/CBIGi001-A-2<br>https://hpscreg.eu/cell-line/CBIGi001-A-3 |
| Ethical/GMO work approvals | McGill University Health Centre Research Ethics Board (DURCAN_IPSC/2019-5374) |
| Addgene/public access repository recombinant DNA sources' disclaimers (if applicable) | pSpCas9(BB)-2A-Puro (PX459) was a gift from Feng Zhang (Addgene plasmid # 48139; http://n2t.net/addgene:48139)<br>RRID: Addgene_48139 |

### Resource utility

PRKN and PINK1 are well known Parkinson’s disease-associated genes. We generated isogenic PRNK-KO, PINK1-KO, and double PINK1-KO/PRKN-KO human iPSC lines, allowing differentiation into neuronal and glial cells for studying Parkinson’s disease and related neurodegenerative disorders. These lines should accelerate understanding of PRKN and PINK1-mediated immune and mitochondrial quality control pathways.

### Resource Details

Two Parkinson’s disease-associated genes, PRKN and PINK1, encode Parkin, a E3-ubiquitin ligase, and the mitochondrial kinase PINK1, proteins critical for mitochondrial quality control (Durcan and Fon, 2015; Sison et al., 2018). We derived PRKN, PINK1 and PINK1/PRKN knockout (KO) lines from AIW002-02 control iPSCs reprogrammed from a 37-year-old healthy male (Chen et al., 2021). Guide RNAs targeting PRKN were cloned into the PX459 vector and transfected into parental AIW002-02 iPSCs. After selection and PCR screening, Sanger sequencing showed one clone contained a homozygous 77 bp deletion in *PRKN* (Fig. 1A), resulting in a frameshift mutation. To knock out PINK1 expression, AIW002-02 or PRKN-KO iPSCs were nuclefected with a combination of CAS9 protein, a sgRNA targeting PINK1 Exon1 (3’ of the ATG start codon), and a ssODN containing a “Stop-tag”. Following limiting dilution, PINK1-KO clones were identified by digital droplet PCR and Sanger sequencing (Fig. 1A), confirming replacement of 17 bp in the PINK1 Exon 1 with a 18 bp sequence containing a stop codons in three reading frames (blue). In the double PINK1-KO/PINK1-KO, however, a homozygous five bp deletion, leading to a frameshift was found (red). Following CRISPR gene KO, qPCR analysis revealed PRKN mRNA expression was < 15% of the expression observed in the parental AIW002-02 iPSCs. In the two PINK1 KO lines, PINK1 expression was also less than 14% of AIW002-02 (Supplementary Fig. 1D). PCR and Sanger sequencing of the top five potential genome-wide off-target sites showed the KO lines lacked off-target mutations or PX459 sequences (Supplementary Fig. 1A, C). Short tandem repeat (STR) profiling demonstrated all KO clones were derived from the AIW002-02 parental line (available from the authors).

**Fig. 1.**
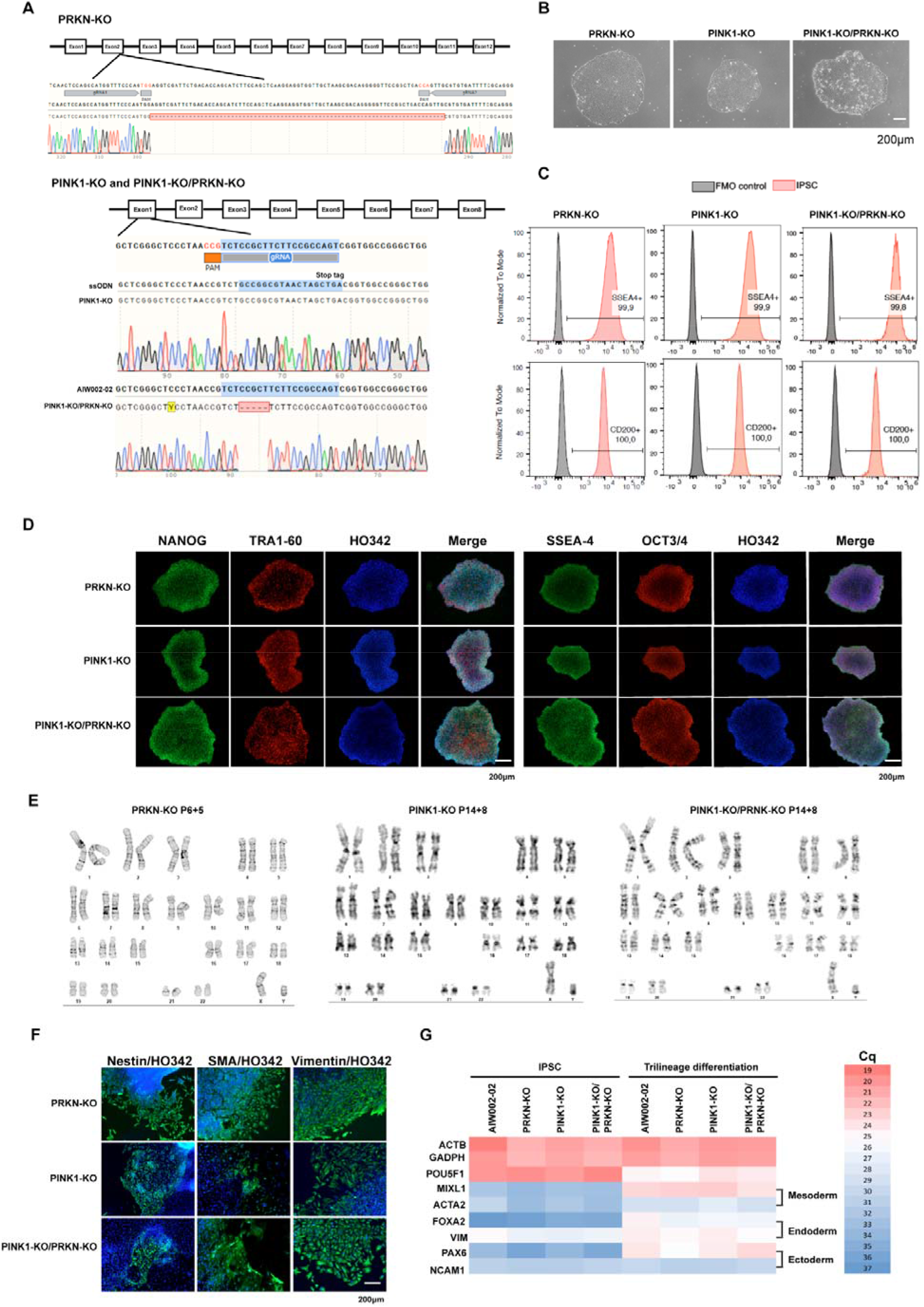
Characterization of isogenic PRKN-KO, PINK1-KO, and PINK1-KO/PRKN-KO iPSC lines.

The PRKN-KO, PINK1-KO and PINK1-KO/PRKN-KO iPSCs showed typical human embryonic stem cell-like morphology by phase-contrast microscopy (Fig. 1B), were ≥99.8% positive for the pluripotency marker SSEA-4 and 100% positive for the iPSC marker CD200 by flow cytometry (Fig. 1C). Immunocytochemistry imaging confirmed all KO iPSCs expressed the pluripotency markers NANOG, TRA1-60, SSEA-4, and OCT3/4 (Fig. 1D). Normal karyotypes were observed for the new KO lines (Fig. 1E) and they did not contain any of the eight most common of karyotypic abnormalities reported in human iPSCs (Supplementary Fig. 1E). All three KO lines were mycoplasma negative as determined by a luminescence-based assay (Supplementary Fig. 1B). Following embryoid body formation and *in vitro* differentiation, immunocytochemistry for ectoderm (nestin), mesoderm (SMA), and endoderm (vimentin) markers demonstrated that PRKN-KO, PINK1-KO and PINK1-KO/PRKN-KO lines were able to form each of the three germ layers. Trilineage differentiation followed by qPCR showed significantly increased expression (lower C_q_ values) of the ectoderm (*PAX6*), endoderm (*FOXA2*) and mesoderm (*ACTA2, MIXL1*) markers (Fig. 1G) compared to iPSCs. In addition, using established protocols (Chen et al., 2019), neuronal precursors from all KO lines were differentiated into tyrosine hydroxylase (TH), MAP2, Tuj1 positive neurons (Supplementary Fig. 1F), indicating that dopaminergic neurons can be generated from these KO iPSCs. These well-characterized isogenic iPSC knockout lines will serve as *in vitro* models for studying PRKN and PINK1-mediated pathways in Parkinson’s disease.

## Materials and Methods

### iPSC culture

AIW002-02 iPSCs were reprogrammed and characterized as described (Chen et al., 2021). iPSCs were maintained in mTeSR1 media (Stemcell Technologies) on Matrigel (Corning)-coated plates at 37 °C with daily media changes, and passaged when 70-80% confluent with Gentle Cell Dissociation Reagent (Stemcell).

### CRISPR/Cas9 knockout

AIW002-02 iPSCs (80,000/well/24-well plate, seeded 24 hours previously) were co-transfected (^~^0.5 μg/vector) using Lipofectomine^™^ Stem Transfection Reagent (ThermoFisher Scientific). Twenty-four hours later, cells were selected in puromycin (0.3ug/ml) for 48h and grown until single clones formed. KO clones were verified by PCR with PRKN-specific primers (Table 2) and Sanger sequencing.

**Table 1:** Characterization and validation

| **Classification**<br>(optional *italicized*) | **Test** | **Result** | **Data** |
| --- | --- | --- | --- |
| **Morphology** | Photography | Normal | Fig. 1 panel B |
| **Pluripotency status evidence for the described cell line** | Qualitative analysis (Immunocytochemistry) | Positive expression of pluripotency markers NANOG, Tra1-60 SSEA-4 and OCT3/4 | Fig. 1 panel D |
|  | Quantitative analysis (Flow cytometry) | PRKN-KO (SSEA-4: 99.9%, CD200: 100%); PINK1-KO (SSEA-4: 99.9%, CD200: 100%); PINK1-KO/PRKN-KO (SSEA-4: 99.8%, CD200: 100%) | Fig. 1 panel C |
| **Karyotype** | Karyotype (G-banding) and resolution | 46,XY<br>Resolution 450–600 | Fig. 1 panel E |
|  | Chromosomal abnormality analysis (qPCR-based) | Normal | Supplementary Fig.1 panel E |
| **Genotyping for the desired genomic alteration/allelic status of the gene of interest** | PCR across the edited site or targeted allele-specific PCR | PCR + sequencing, homozygous | Fig. 1 panel A |
|  | Transgene-specific PCR | N/A | N/A |
| **Verification of the absence of random plasmid integration events** | PCR | Absence of pSpCas9(BB)-2A-Puro (PX459) integration in PRKN-KO | Supplementary Fig.1 panel C |
| **Parental and modified cell line genetic identity evidence** | STR analysis | 10 specific loci were tested. All isogenic lines matched 100% | Supplementary data 1<br>Submitted in archive with journal |
| **Mutagenesis / genetic modification outcome analysis** | Sequencing (genomic DNA PCR or RT-PCR product) | PRKN-KO: Homozygous, 77 bp deletion in *PRKN*<br>PINK1-KO: Homozygous, 18 bp indel in *PINK1*<br>PINK1-KO/PRKN-KO: Homozygous, 5 bp deletion in *PINK1* | Fig. 1 panel A |
|  | PCR-based analyses (qPCR) | Decreased expression of PRKN and/or PINK1 in KO lines | Supplementary Fig.1 panel D |
|  | Southern Blot or WGS; western blotting (for knock-outs, KOs) | Not performed | N/A |
| **Off-target nuclease analysis** | PCR across top 5 predicted top likely off-target sites | Lack of NHEJ-caused mutagenesis in the top predicted off-target Cas nuclease activity | Supplementary Fig.1 panel A |
| **Specific pathogen-free status** | Mycoplasma | Micoplasma testing by luminescence. Negative | Supplementary Fig. 1 panel B |
| **Multilineage differentiation potential** | Embryoid body formation (Immunocytochemistry) | Proof of differentiation into three germ layers by immunostaining for Nestin , SMA, and Vimentin | Fig. 1 panel F |
|  | Directed differentiation mRNA expression (qPCR) | Increased expression of *PAX6* (ectoderm ), *MIXL1* (mesoderm), *FOXA2 and Vimentin* (endoderm) | Fig. 1 panel G |
|  | Directed differentiation (Immunocytochemistry) | Dopaminergic neuron markers (TH, MAP2, Tuj) | Supplementary Fig.1 panel F |
| **Donor screening (OPTIONAL)** | HIV 1 + 2 Hepatitis B, Hepatitis C | Negative | Available from the authors |
| **Genotype - additional histocompatibility info (OPTIONAL)** | Blood group genotyping | Not performed | N/A |
|  | HLA tissue typing | Not performed | N/A |

**Table 2:**
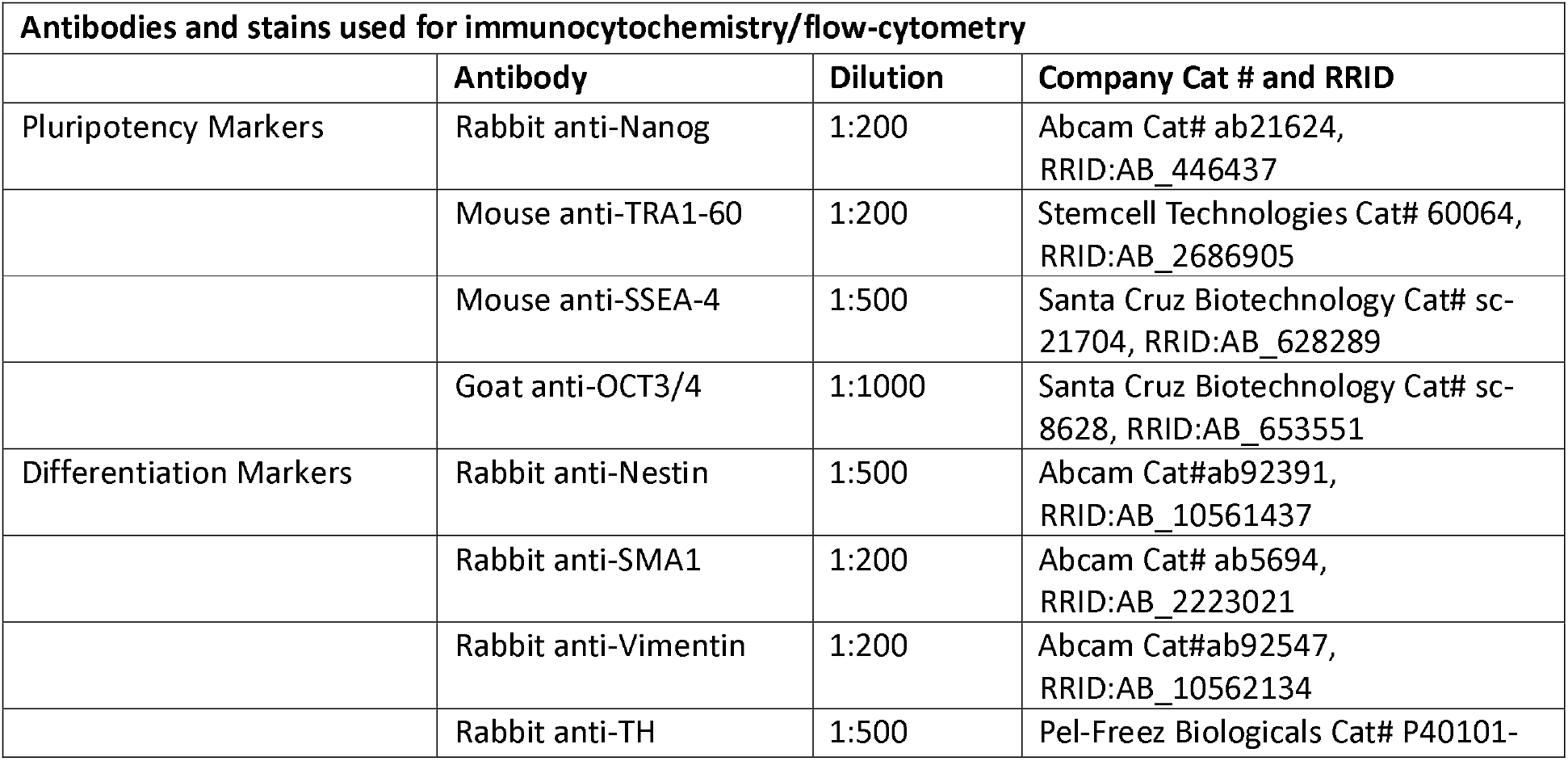
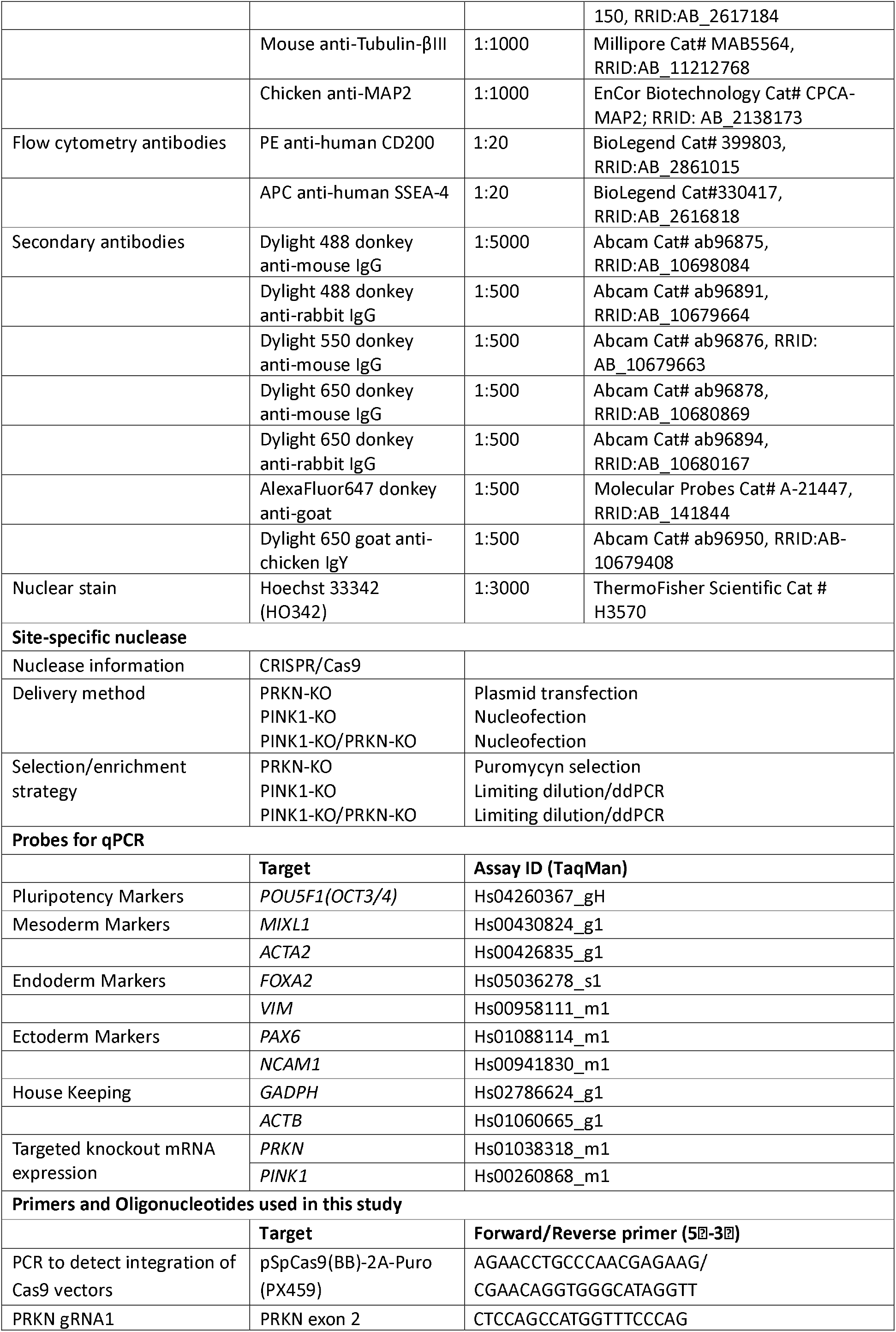
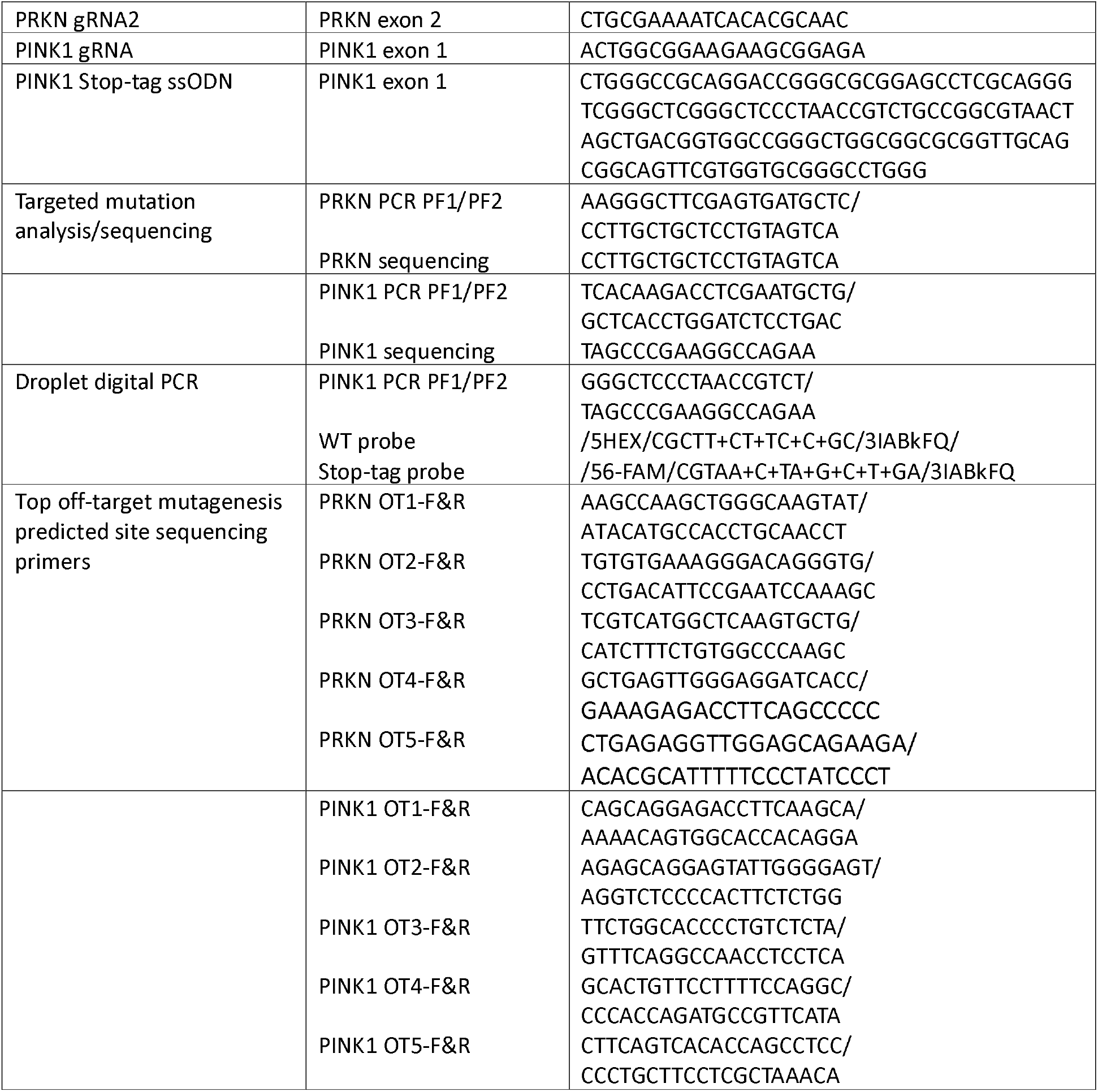
Reagents details

Alternatively, ribonucleoprotein complexes containing CAS9 protein (1 μl; stock 61 μM), PINK1 sgRNA (3 μl; stock 100 μM) and ssODN (1 μl,stock 100 μM) in 20 μl of nucleofection buffer P3 were nucleofected (program CA137, 4D-Nucleofector Device, Lonza) into 500,000 detached iPSCs (Deneault et al., 2021). After limiting dilution, KO clones were identified by ddPCR (QX200™ Droplet Reader, Bio-Rad) and Sanger sequencing.

### Flow cytometry

iPSCs were detached using TrypLE Express (ThermoFisher Scientific), stained with Live Dead Aqua (Invitrogen) for viability, then blocked with Human TruStain FcX (Biolegend). Cells were stained with FACS antibodes (Table 2) for 30 min at RT, before data acquisition on an Attune NxT Flow Cytometer (ThermoFisher Scientific) and FlowJo analysis.

### Immunocytochemistry

Cells were fixed in 4% paraformaldehyde at RT for 20 mins, permeabilized with 0.2% Triton X-100/PBS for 10 min, blocked in 5% donkey serum/1% BSA/0.05% Triton X-100/PBS for 2 h, incubated with primary antibodies (Table 2) in blocking buffer overnight at 4°C, secondary antibodies (Table 2) for 2 h at RT, then counterstained with Hoechst 33342 for 5 min. Images were acquired using the automated Evos FL-Auto2 imaging system (ThermoFisher Scientific).

### Karyotyping and genomic integrity analysis

G-band karyotyping was performed by Wicell Cytogenetics, Madison, WI. Genomic integrity was analyzed using DNA extracted from iPSCs with a Genomic DNA Mini Kit (Geneaid) and the hPSC Genetic Analysis Kit (Stemcell).

### Three Germ Layer Differentiation

Embyoid bodies (EBs) were generated from iPSCs, allowed to spontaneously differentiate, fixed, and analyzed by immunocytochemistry. Pluripotency was also assessed with The STEMdiff Trilineage differentiation kit (Stemcell) (Chen et al., 2021).

### qPCR analysis

cDNA was generated (iScript Reverse Transcription Supermix; BioRad) from purified RNA (NucleoSpin RNA kit; Takara). Raw data from the QuantStudio 5 Real-Time PCR System (Applied Biosystems) were processed with the Auto-qPCR app using GADPH or ACTB expression as endogenous controls (Chen et al., 2021).

### Dopaminergic neuron differentiation

iPSCs were induced into DA-NPCs according to established protocols (Chen et al., 2019). DA-NPCs were differentiated into neurons in neurobasal medium, supplemented with N2 and B27, 1 μg/ml laminin, 500 μM db-cAMP, 20 ng/ml BDNF, 20 ng/ml GDNF, 200 μM ascorbic acid, 100 nM Compound A and 1 ng/ml TGF-β. Immunostaining was conducted at 4 weeks maturation.

### Short Tandem Repeat Analysis

STR analysis was performed with the GenePrint® 10 System (Promega) at The Centre for Applied Genomics, The Hospital for Sick Children, Toronto, ON.

### Mycoplasma

iPSC cuture media were tested using the luminescence-based MycoAlert Mycoplasma Detection kit (Lonza).

## Acknowledgements

TMD received funding to support this project from the Canada First Research Excellence Fund, awarded through the Healthy Brains, Healthy Lives initiative at McGill University, the CQDM FACs program, the Michael J. Fox Foundation and a project grant from CIHR (PJT – 169095).

## Appendix A. Supplementary data

**Supplementary Fig. 1.**
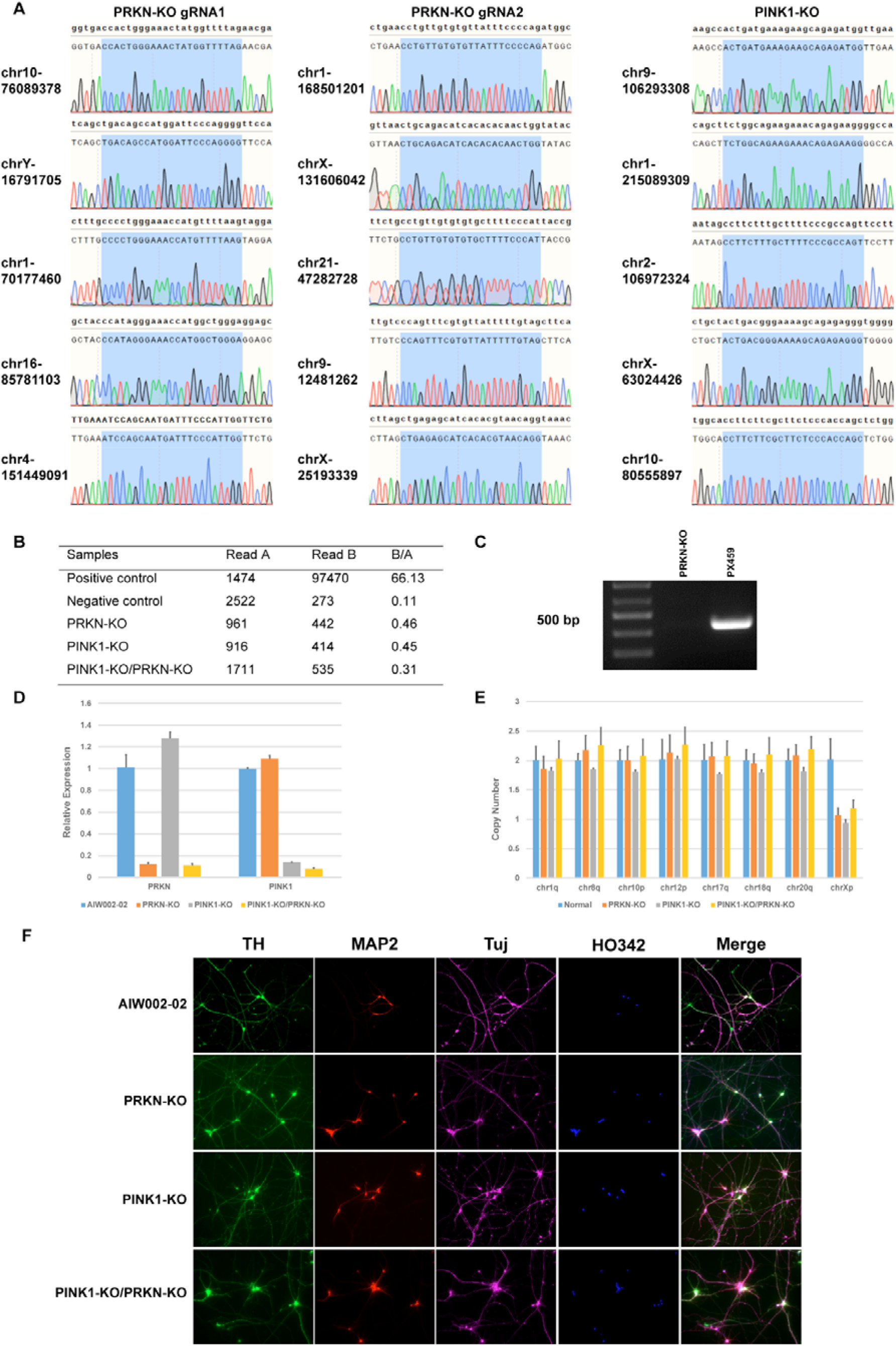

## References

Chen, C.X.-Q., Abdian, N., Maussion, G., Thomas, R.A., Demirova, I., Cai, E., Tabatabaei, M., Beitel, L.K., Karamchandani, J., Fon, E.A., Durcan, T.M., 2021. A multistep workflow to evaluate newly generated iPSCs and their ability to generate different cell types. Methods Protoc. 4(3), 50. https://www.mdpi.com/2409-9279/4/3/50.

Chen, C.X.-Q., Lauinger, N., Rocha, C., Rao, T., Durcan, T.M., 2019. Generation of dopaminergic or cortical neurons from neuronal progenitors. Zenodo. https://doi.org/10.5281/zenodo.3738323.

Deneault, E., Chaineau, M., Nicouleau, M., Castellanos Montiel, M.J., Franco Flores, A.K., Haghi, G., Chen, C.X.-Q., Abdian, N., Shlaifer, I., Beitel, L.K., Durcan, T.M., 2021. A streamlined CRISPR workflow to introduce mutations and generate isogenic iPSCs for modeling amyotrophic lateral sclerosis. Methods. https://doi.org/10.1016/j.ymeth.2021.09.002.

Durcan, T.M., Fon, E.A., 2015. The three ‘P’s of mitophagy: PARKIN, PINK1, and post-translational modifications. Genes Dev. 29(10), 989–999. http://www.genesdev.org/cgi/doi/10.1101/gad.262758.115.

Sison, S.L., Vermilyea, S.C., Emborg, M.E., Ebert, A.D., 2018. Using patient-derived induced pluripotent stem cells to identify Parkinson’s disease-relevant phenotypes. Curr. Neurol. Neurosci. Rep. 18(12), 84. https://www.ncbi.nlm.nih.gov/pubmed/30284665.

